# Bi-directional control of a prelimbic somatostatin microcircuit decreases binge alcohol consumption

**DOI:** 10.1101/2020.11.27.400465

**Authors:** Nigel C. Dao, Dakota F. Brockway, Malini Suresh Nair, Nicole A. Crowley

**Author notes:** Correspondence: Nicole A. Crowley, 326 Mueller Lab, Penn State University, University Park PA 16802.

## Abstract

Somatostatin neurons have been implicated in a variety of neuropsychiatric disorders such as depression and anxiety, but their role in substance abuse disorders, including alcohol use disorder (AUD), is not fully characterized. Here we found that repeat cycles of alcohol binge drinking in the Drinking-in-the-Dark (DID) model led to hypoactivity of somatostatin (SST) neuronal in the prelimbic (PL) cortex by diminishing their action potential firing capacity and excitatory/inhibitory transmission dynamic. We examined their role in regulating alcohol consumption via bidirectional chemogenetic manipulation. Both hM3Dq-induced excitation and KORD-induced silencing of PL SST neurons paradoxically reduced alcohol binge drinking in males and females, with no effect on sucrose consumption. This effect is mediated directly via monosynaptic connection from SST neurons onto pyramidal neurons and indirectly via an intermediate GABAergic source. Optogenetic-assisted circuit mapping revealed that PL SST neurons preferentially synapse onto pyramidal neurons over other GABAergic populations in males, whereas SST neuron-mediated inhibition is balanced across cell types in females. Alcohol binge drinking disinhibits pyramidal neurons by augmenting SST neurons-mediated GABA release and synaptic strength onto other GABAergic populations. Together these data suggest substantial interaction between alcohol binge drinking and SST neurons inhibitory circuit in the PL, as well as provide evidence for these neurons as a potential therapeutic candidate for the treatment of alcohol use disorders, including binge drinking.

## INTRODUCTION

Alcohol use disorder (AUD) is a chronically relapsing disorder (with three sub-classifications: mild, moderate, severe), and characterized by the presence of a minimum of 2 symptoms related to both alcohol consumption behavior and the ramifications of that behavior (1,2). Binge drinking, in particular, has been shown to have severe negative health outcomes, with over 26% of adults (18+) reporting binge drinking in the past month (3). The National Institute on Alcohol Abuse and Alcoholism (NIAAA) defines binge drinking as a consumption pattern leading to a blood alcohol content greater than 80 mg/dL in a 2 hr period (approximately 4 drinks for women or 5 drinks for men). Though both men and women engage in binge drinking, sex differences exist in both onset and patterns of binge drinking and outcomes (for overview, see (4).

Multiple brain regions, neuronal subtypes, neurotransmitters, and neuromodulators have been implicated in alcohol’s effects (5). The prelimbic (PL) cortex is a key brain region in addiction and substance abuse (6,7), with projections to many limbic areas (7) including the bed nucleus of the stria terminalis (8,9) and the nucleus accumbens (10) to provide top-down regulation of affective and motivational behaviors. It receives excitatory glutamate afferents from the basolateral amygdala (11,12), and the hippocampus (13), and inhibitory control from a variety of subpopulations of gamma aminobutyric acide (GABA) neurons that express peptidergic markers and receptors such as dynorphin (DYN; 14), corticotropin-releasing factor (CRF; (15,16)), neuropeptide-y (NPY;(17)) and somatostatin (SST; (18,19)). Neurotransmission within the PL is known to be altered in multiple animal models of alcohol exposures, including binge drinking (20) and vapor ethanol exposure (21,22). Markedly, recent work has demonstrated that repeated cycles of binge drinking diminishes NPY expression in the PL, and that activation of NPY signaling within the PL otherwise can reduce binge drinking (17), highlighting the engagement of these peptide-expressing GABA neurons in AUD and their therapeutic potentials.

Human and preclinical rodent evidence points to SST neurons as a key modulator of neuropsychiatric disorders, in particular anxiety and depression (23). Global genetic upregulation of SST neurons reduces anxiety-like and depression-like behavior in mice (24), with site-specific manipulations of PL SST neurons having a similar effect (25). In addition, PL SST neurons specifically encode fear-related memories in fear conditioning (26)and social fear (27). Despite the strong implications for SST neurons as a positive target for mood disorders, its role in AUD remains to be explored. Here, we demonstrate that SST neurons are strongly modulated by binge alcohol consumption in both male and female mice, and that bi-directional chemogenetic modulation of SST neurons in the PL reduces binge alcohol consumption in both sexes (with a greater overall effect in females). In addition, we provide new insight into the overall circuitry of SST neurons within the PL and SST neurons’ role maintaining signaling of PL pyramidal output neurons. This work suggests SST neurons, specifically in the PL, may serve as a promising therapeutic target for AUD.

## METHODS

### Animals

All experiments were approved by the Pennsylvania State University Institutional Animal Care and Use Committee. Adult male and female C57BL/6J (stock #000664, The Jackson Laboratory), hemizygous SST-IRES-Cre mice (stock #013044, The Jackson Laboratory) and *Ai9* reporter mice (stock #007909, The Jackson Laboratory) on C57BL6/J background were bred in-house and genotyped by standard PCR protocol. All mice were single-housed on a 12 hr reverse light cycle for the entirety of the experiments (lights off at 7:00am), as is consistent with choice alcohol consumption paradigms (28,29). Mice had *ad libitum* access to food and water (except for during the DID procedure, described below, when water was removed).

### Surgeries

At 8 weeks of age, mice underwent intra-PL (from Bregma, AP: +1.8mm; ML: ±0.4mm, DV: −2.3mm) viral injections. Mice were deeply anesthetized with isoflurane (5% induction, 1-2% maintenance) and mounted on the stereotaxic frame (Stoelting, Wood Dale, IL). Following craniotomy, 0.3 ul of virus per side was injected into the PL at a rate of 0.1 ul/min via a 1ul Hamilton Neuros Syringe. The syringe was left in place for an extra 5 minutes before being slowly removed to limit efflux of virus. Bupivacaine (4 mg/kg) and ketoprofen (5 mg/kg) were applied topically and intraperitoneally, respectively, for postoperative pain management. Following surgery, mice were monitored and allowed to recover for one week. At least 3 weeks elapsed between the time of viral injection and experimental manipulations (Designer Receptors Exclusively Activated by Designer Drugs, DREADD; ligands administration or electrophysiology) to allow for ample viral expression.

### Viral Vectors

Viral vectors AAV5-hSyn-DIO-mCherry, AAV8-hSyn-DIO-hM3D(Gq)-mCherry, AAV8-hSyn-dF-HA-KORD-IRES-mCitrine, AAV5-EF1a-dF-hChR2(H134R)-eYFP-WPRE-HGHpA, AAV5-CaMKIIa-hChR2(H134R)-EYFP, and AAV5-CaMKIIa-hM4D(Gi)-mCherry, described elsewhere (30,31,32), were obtained from Addgene (Watertown, MA).

### Drinking in the Dark (DID)

DID was conducted as previously published (20,29). Mice received 20% (v/v) ethanol (EtOH; Koptec, Decon Labs, King of Prussia, PA) for 2 hr in drinking water, 3 hr into the dark cycle (i.e. 10 a.m.) on three sequential days. On the fourth day, they received 20% ethanol for 4 hr. Following the binge day, mice had three days of abstinence before repeating the cycle (number of cycles indicated per experiment below). When applicable, mice underwent a baseline week of DID before surgeries.

### Sucrose DID

In some cohorts, four days after the last EtOH binge session, mice underwent sucrose DID conducted as previously published (17,33). Mice received 10% sucrose in water 3 hr into the dark cycle on three sequential days. On the fourth day, they received 10% (w/v) sucrose in tap water for 4 hr. The mice underwent three cycles total.

### Drug Administration

Clozapine *n*-oxide (CNO; #HB1807, HelloBio, Princeton, NJ) was dissolved to 30 mg/ml in pure DMSO (Fisher Scientific, Waltham, MA) and subsequently diluted to 0.3 mg/ml in 0.9% saline. Salvinorin B (SalB; #HB4887, HelloBio, Princeton, NJ) was dissolved to 10 mg/ml in DMSO. Mice were habituated to handling for injections for three consecutive days before the first vehicle injection. For EtOH DID, mice received vehicle injections of either 1% DMSO in saline (10 ml/kg; intraperitoneal, i.p.) or pure DMSO (1 ml/kg; subcutaneously, s.c.) on cycle 2. Injections of CNO and SalB were counterbalanced across mice to account for any order effects (i.e., mice received either CNO (3 mg/kg; i.p.) or SalB (10 mg/kg; s.c.) on cycle 3, and the other drug on cycle 4). For sucrose DID, mice received vehicle injections on cycle 1, and CNO or SalB injections on cycle 2 or 3. For hM4D-mediated inhibition of pyramidal neurons experiment, mice were administered saline on cycle 2 and CNO (3 mg/kg) on cycle 3. All injections were done 30-40 minutes before the start of the binge session (30).

### Electrophysiology

Whole cell voltage clamp and current clamp recordings were conducted in the PL similarly to those previously published (spontaneous excitatory and inhibitory postsynaptic currents, sEPSCs and sIPSCs respectivelu, and intrinsic excitability(20,21,34); confirmation of DREADD expression and function (30)). The PL was identified according to the Allen Mouse Brain Atlas. Mice were deeply anesthetized via inhaled isoflurane and rapidly decapitated. Brains were rapidly removed and processed according to the NMDG protective recovery method (35). Briefly, brains were immediately placed in ice-cold oxygenated N-methyl-D-glucamine (NMDG)-HEPES aCSF containing the following, in mM: 92 NMDG, 2.5 KCl, 1.25 NaH2PO4, 30 NaHCO3, 20 HEPES, 25 glucose, 2 thiourea, 5 Na-ascorbate, 3 Na-pyruvate, 0.5 CaCl2·2H2O, and 10 MgSO4·7H2O (pH to 7.3–7.4). 300μM coronal slices containing the PL were prepared on a Compresstome vibrating microtome VF-300-0Z (Precisionary Instruments, Greenville, NC), and transferred to heated (31°C) NMDG-HEPES aCSF for a maximum of 10 min. Slices were then transferred to heated (31°C) oxygenated normal aCSF (in mM: 124 NaCl, 4.4 KCl, 2 CaCl2, 1.2 MgSO4, 1 NaH2PO4, 10.0 glucose, and 26.0 NaHCO3, pH 7.4, mOsm 300-310), where they were allowed to rest for at least 1h before use. Finally, slices were moved to a submerged recording chamber (Warner Instruments, Hamden, CT) where they were continuously perfused with the recording aCSF at room temperature. Recording electrodes (3–6 MΩ) were pulled from thin-walled borosilicate glass capillaries with a Narishige P-100 Puller.

Pyramidal neurons in layer 2/3 of the PL cortex were identified by location, morphology (prominent triangular soma and apical dendrites), and membrane characteristics (high capacitance > 75 pF, or low membrane resistance < 100 mΩ). SST-expressing neurons were identified in SST-IRES-Cre::Ai9 mice via presence of tdTomato flourescence under a 40x immersed objective with 565 nm LED excitation. Non-SST expressing neurons were identified by lack of fluorescence, morphology and membrane characteristics (low capacitance, and high membrane resistance > 200 mΩ).

Spontaneous AMPA receptor-mediated excitatory postsynaptic currents (sEPSCs) and GABA_A_ receptor-mediated inhibitory postsynaptic currents (sIPSCs) were measured in voltage-clamp (−55 mV and +10 mV, respectively) using electrodes filled with a cesium-methanesulfonate (Cs-Meth) intracellular recording solution (containing the following, in mM: 135 Cs-methanesulfonate, 10 KCl, 10 HEPES, 1 MgCl2, 0.2 EGTA, 4 Mg-ATP, 0.3 GTP, 20 phosphocreatine, 287-290 mOsm, pH 7.33). Frequency (Hz) and amplitude (pA) of sIPSCs within individual neurons were measured in a 2-minute epoch following 10 minutes of stabilization. Measurements of intrinsic excitability were recorded in current clamp, including resting membrane potential (RMP), rheobase (the minimum amount of current needed to elicit an action potential), action potential threshold (the membrane potential at which the first action potential fired), and current-injection induced firing (0-200 pA, 10 pA per step). Experiments were performed at both RMP and at the holding potential −70 mV. Electrodes were filled with a potassium gluconate-based (KGluc) intracellular recording solution (in mM: 135 K-Gluc, 5 NaCl, 2 MgCl2, 10 HEPES, 0.6 EGTA, 4 Na2ATP, and 0.4 Na2GTP, 287-290 mOsm, pH 7.35).

In separate experiments, to confirm the functionality of the hM3Dq and KORD viruses, CNO (10 μM) or SalB (1 μM) was added to the recording aCSF for 10 min following a 10-min stabilization of RMP and recording continued for a 15-min washout. Tetrodotoxin (500 nM) was added to the recording aCSF to block action potentials for consistent measurement of RMP. Average RMP were normalized to a 5-min baseline. To record sIPSCs on pyramidal neurons during CNO and SalB bath application, 3 mM kynurenic acid was included in the recording aCSF to block α-amino-3-hydroxy-5-methyl-4-isoxazolepropionic acid (AMPA) and N-methyl-D-aspartate (NMDA) receptor-mediated excitatory postsynaptic currents (EPSCs), using a potassium chloride/ potassium-gluconate (KCl-KGluc) intracellular recording solution (in mM: 70 potassium gluconate, 80 KCl, 10 HEPES, 1 EGTA, 4 Na2ATP, 0.4 Na2GTP, 287-290 mOsm, pH 7.3). Neurons were held at −70 mV. After a stable 6-min baseline, CNO (10 μM) or SalB (1 μM) in aCSF+kynurenic acid was perfused onto the slice for 10 min, followed by 15 min of aCSF+kynurenic acid only washout. Frequency and amplitude of postsynaptic events were normalized to a 6-min baseline period.

To assess the local connectivity between SST neurons and both pyramidal neurons and non-SST putative GABAergic neurons, optogenetically-induced light evoked IPSCs were elicited by 2 x 1-ms pulses of 470 nM blue light delivered 100 ms apart (Cool LED, Traverse City, MI, USA), using the K-gluc-based intracellular recording solution (see above). Neurons were held at −50 mV, and photostimulation of SST neurons produced outward currents onto pyramidal neurons and non-SST neurons that are sensitive to blockage by the GABA_A_ receptor antagonist picrotoxin (100 μM; Supplementary Figure 2A). Evoked IPSC peak amplitude induced by the first LED pulse and paired-pulse ratio (PPR: amplitude of pulse 2/amplitude of pulse 1) were measured as proxies of synaptic strength between SST neurons and other neuronal populations in the PL cortex. Photostimulation were performed across multiple laser intensities (10%-100%, 10% interval), however the peak optically-evoked IPSC amplitude did not significantly vary with laser intensity so only data derived from maximum laser intensity are reported.

Signals were digitized at 10 kHz and filtered at 3 kHz using a Multiclamp 700B amplifier and analyzed using Clampfit 10.7 software (Molecular Devices, Sunnyvale, CA). For all measures, recordings were performed in a maximum of two neurons per subregion, per mouse, and lowercase n values reported reflect the number of neurons for each measure.

### Histology

Mice were deeply anesthetized with Avertin (250 mg/kg), and transcardially perfused with ice-cold phosphate buffered saline (PBS) then 4% (w/v) paraformaldehyde (PFA). Brains were post-fixed in PFA overnight, and sectioned at 40 microns using a Compresstome vibrating microtome. PL cortex-containing sections were incubated in 1:20,000 DAPI in PBS for 30 min, mounted on SuperFrost glass slides, air-dried, and coverslipped with ImmunoMount mounting media. Viral injections were assessed under mCherry and/or mCitrine fluorescence filter in an Olympus BX63 upright microscope (Center Valley, PA), and the brightest fluorescence point was chosen as the center of the injection. Mice with unilateral viral expression, and/or missed injections (including expression in the infralimbic cortex) were excluded from analysis. Quantification of overall injection localization was performed in ImageJ (NIH) and presented in **Figure 2B** and **Supplementary Figure 2**. Cells visualizable with both mCherry and mCitrine fluorescence were counted as mCitrine+/mCherry+, whereas those visualizable with either mCherry or mCitrine only were counted as mCherry+ or mCitrine+, respectively.

### Statistical Analysis

Experimenters were blinded to group (alcohol or water), and chemogenetic manipulation whenever possible. Data was analyzed by ANOVAs and post-hoc test as appropriate and indicated for each experiment. Statistical significance threshold was set at α = 0.05. Statistical analysis and graphs were performed in Graphpad Prism 7.0, and finalized figures were constructed in Adobe Illustrator and BioRender. Data presented show Means and Standard Error of the Mean (SEM) where applicable.

## RESULTS

### Alcohol binge-like drinking alters PL cortex SST neuronal intrinsic excitability and dampens inhibitory inputs onto pyramidal neurons

To examine DID alcohol consumption-induced neuroalterations in PL SST neurons, *SST*-*IRES-CRE::Ai9* male and female mice underwent DID for four cycles (**Figure 1A** for representative expression, **1B** for experimental timeline). Similar to previous reports, female mice consumed higher levels of alcohol than male mice (**Supplementary Figure 1**). 24 hr after the last alcohol binge drinking session, we performed whole-cell patch clamp electrophysiology in PL SST neurons to examine DID-induced alterations in intrinsic excitability in and synaptic transmission on these neurons. First, current clamp experiments revealed that PL SST neurons exhibited marked changes in intrinsic excitability. Male SST neurons showed similar resting membrane potential (RMP), rheobase and action potential threshold at −70mV relative to controls (**Figure 1C, D, E**), but reduced firing during the voltage-current steps at −70mV (**Figure 1F**). In females, SST neurons displayed depolarized RMP compared to controls (**Figure 1C**). Interestingly, when held at an RMP of −70 mV, these neurons showed dampened excitability, with a significantly higher rheobase (**Figure 1E**), reduced number of action potentials fired during a voltage-current plot protocol (**Figure 1G**) and no changes in action potential threshold (**Figure 1D**). Additionally, voltage clamp experiments revealed that alcohol binge drinking disrupted the excitation/inhibition dynamics in PL SST neurons. Both male and female SST neurons showed an attenuation of sEPSC frequency (**Figure 1H, I)**, with no changes in sEPSC amplitude (**Figure 1J).** While alcohol binge drinking did not impact sIPSC frequency across sexes and drinking groups (**Figure 1K**), it produced opposite adaptations in PL SST neuronal sIPSC amplitude between sexes (**Figure 1L)**. Whereas alcohol binge drinking reduced sIPSC amplitude in male PL SST neurons, it otherwise augmented sIPSC amplitude in female SST neurons. Notably, there is also a basal sex difference in control mice, where male SST neurons display larger sIPSC events than female SST neurons, suggesting significant variability in the postsynaptic expression of GABA_A_ receptor between sexes that could be differentially altered by a history of binge drinking.Together, these data demonstrated that repeated cycles of alcohol binge drinking result in a state of hypoexcitability in PL SST neurons via a reduction in intrinsic excitability and excitatory synaptic transmission, which could ultimately lead to disinhibition of neighboring populations.

**Figure 1:**
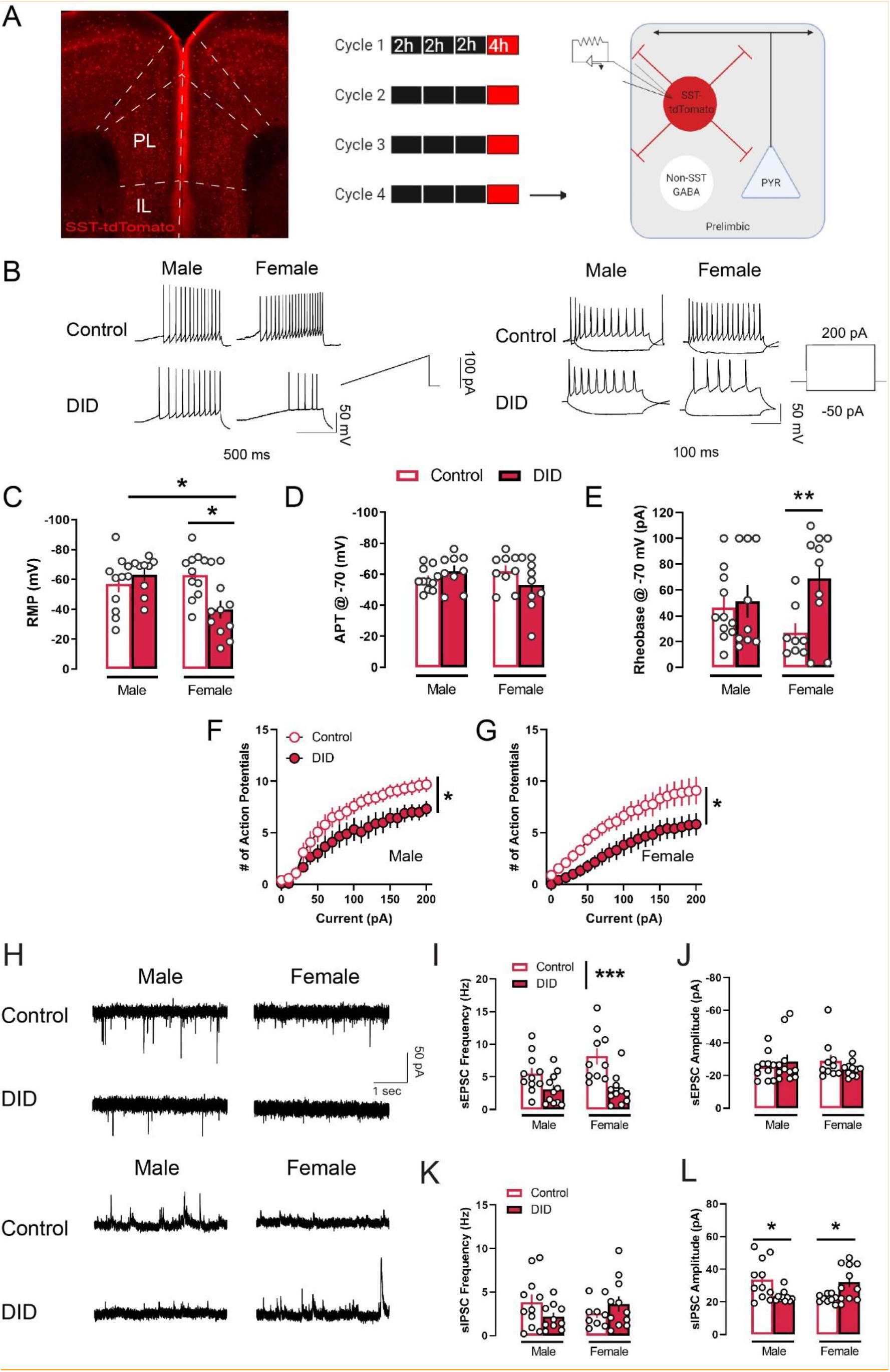
Binge-like alcohol consumption dampens intrinsic excitability and excitatory drive of SST neurons in the PL. (A) (Left) PL SST neurons under tdTomato fluorescence. (Right) Schematic of experimental timeline. (B) Representative traces of rheobase and VI recording in SST cells in the PL. (C) Binge drinking depolarizes resting membrane potential in female SST PL neurons (DID: n = 11 cells; control: n = 11 cells, control female vs. DID female: *p* = 0.014, DID male vs. DID female: *p* = 0.019), but not male (DID: n = 9 cells; control: n = 11 cells; F_sex_(1,38) = 2.752, *p* = 0.105; F_binge_(1,38) = 2.539, *p* = 0.119, F_sex x binge_(1,38) = 7.896, *p* = 0.007). (D) Action potential threshold in SST PL neurons across sexes was not altered following binge drinking (F_sex_(1,34) = 1.107, *p* = 0.3; F_binge_(1,34) = 0.017, *p* = 0.894, F_sex x binge_(1,34) = 1.336, *p* = 0.256) (E) Binge drinking increases the minimum current needed to elicit an action potential in female SST PL neurons (DID: n = 10 cells, control: n = 8 cells, control female vs. DID female: *p* = 0.018), but not males (DID: n = 9 cells, control: n = 11 cells; F_sex_(1,34) = 0.0, *p* = 0.941; F_binge_(1,34) = 4.967, *p* = 0.032, F_sex x binge_(1,34) = 3.097, *p* = 0.087). (F) Current-induced spiking was reduced in male SST PL neurons following binge drinking (DID: n = 9 cells, control: n = 10 cells; F_current_(20,340) = 71.079, *p* < 0.001; F_binge_(1,17) = 3.887, *p* = 0.061, F_current x binge_(20,340) = 1.734, *p* = 0.026). (G) Binge drinking reduces current-induced spiking in female SST PL neurons (DID: n = 12 cells, control: n = 9 cells, F_current_(20,380) = 39.339, *p* < 0.001; F_binge_(1,19) = 5.754, *p* = 0.027, F_current x binge_(20,380) = 0.995, *p* = 0.467) (H) Representative sEPSC traces (top four) and sIPSC traces (bottom four) in SST cells in the PL. (I) Binge drinking reduced sEPSC frequency in SST cells in both sexes (control male: n = 11 cells, DID male: n = 11 cells, control female: n = 10 cells, DID female: n = 11 cells, F_sex_(1,39) = 2.101, *p* = 0.155; F_binge_(1,39) = 18.649, *p* < 0.001, F_sex x binge_(1,39) = 2.472, *p* = 0.123). (J) Binge drinking did not alter sEPSC amplitude in SST cells (F_sex_(1,38) = 0.024, *p* = 0.8763; F_binge_(1,39) = 0.066, *p* = 0.798, F_sex x binge_(1,39) = 1.704, *p* = 0.199) (K) Binge drinking did not alter sIPSC frequency in SST cells (control male: n = 11 cells, DID male: n = 10 cells, control female: n = 9 cells, DID female: n = 11 cells; F_sex_(1,37) = 0.021, *p* = 0.884; F_binge_(1,37) = 0.152, *p* = 0.698, F_sex x binge_(1,37) = 3.232, *p* = 0.080). (L) sIPSC amplitude in SST cells exhibit basal sex differences, in which male SST neurons have larger inhibitory current amplitude than female SST neurons (*p* = 0.014). Binge drinking sex-dependently altered sIPSC amplitude in SST cells, in which sIPSC amplitude was reduced in males (*p* = 0.018), but increased in females (*p* = 0.029) (F_sex_(1,37) = 0.25, *p* = 0.62; F_binge_(1,37) = 0.007, *p* = 0.932, F_sex x binge_(1,37) = 14.13, *p* < 0.001). DID: N = 6 male mice and 5 female mice, control: N = 5 male mice and 5 female mice. * *p* < 0.05. ** *p* < 0.01. *** *p* < 0.001. Bonferroni’s post-hoc test.

### Both chemogenetic activation and inhibition of SST neuron activity reduce binge-like alcohol drinking via direct and indirect inhibition of pyramidal neurons

Given previous reports on the resiliency-conferring role of PL SST neurons in affective disorders (23,24) and our observation of a reduction in both SST neuronal excitability and complementary inhibitory transmission onto PL pyramidal neurons post-DID, we investigated the ability of chemogenetic manipulation of PL SST neurons to alter drinking. We injected SST-IRES-Cre male and female mice with a cocktail of AAVs encoding the excitatory Gq-coupled hM3Dq and the inhibitory Gi-coupled kappa-opioid-derived DREADD (KORD) into the PL, which allowed for bidirectional manipulation of the same SST neurons during DID (26) (for experimental timeline, see **Figure 2A,** representative viral injections, **Figure 2B;** see **Supplementary Figure 1** for quantification of overlap between hM3Dq and KORD).

**Figure 2:**
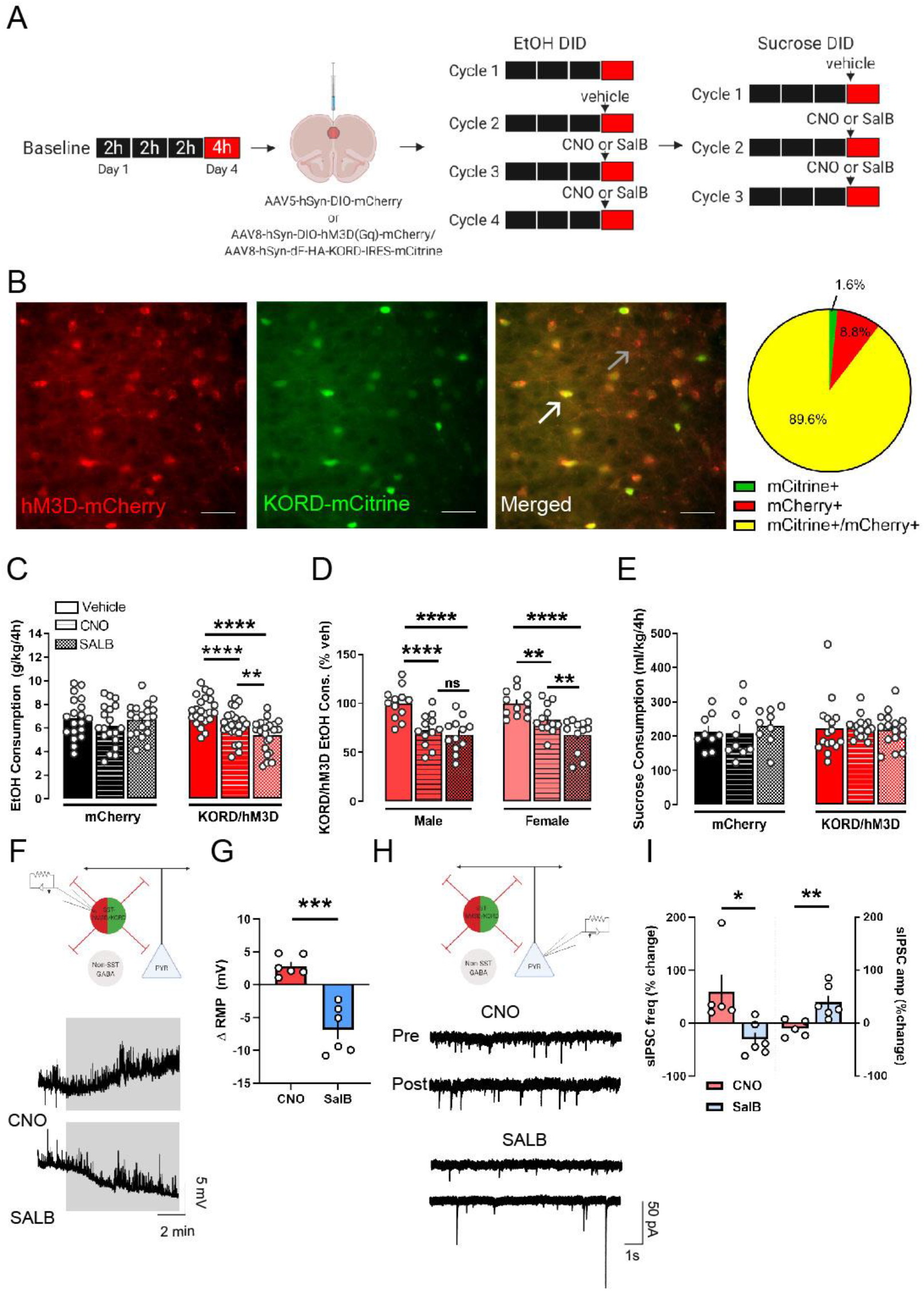
Bi-directional control of PL SST neurons reduces binge drinking. (A) Schematic of the experimental design. Both male and female SST-IRES-Cre mice were used in this experiment. (B) Representative images of hM3Dq- and KORD-expressing SST neurons at high magnification (40x). White arrows: mCitrine+/mCherry+ cells. Black arrows: mCherry+ only cells. Scale bar 50 μM. (Right) Percent of total infected SST neurons in the PL expressing mCitrine-tagged KORD (green) only, mCherry-tagged hM3Dq only (red), or both viruses (yellow). (C) CNO-induced activation and SalB-induced silencing of SST PL neurons reduce binge drinking in hM3Dq- and KORD-expressing mice (N = 23), but not in mCherry-expressing control mice (N = 19) (F_drug_(2,80) = 17.1, *p* < 0.00001, F_virus_(1, 40) = 0.289, *p* = 0.593, F_virus x drug_(2,80) = 11.79, *p* < 0.0001). Additionally, silencing of SST PL neurons might be more effect in dampening binge drinking than activation of the same neuronal population (CNO vs. SalB, *p* = 0.009). Data included both males and females. (D) In hM3Dq- and KORD-expressing mice, males (N = 12) are equally susceptible to activation and silencing of SST PL neurons (CNO vs. SalB, *p* = 0.70), whereas activation of SST PL neurons in females (N = 12) might be less effective (CNO vs. SalB, *p* = 0.009) (F_drug_(1,22) = 0.484, *p* = 0.494, F_sex_(2,44) = 49.08, *p* < 0.001, F_sex x drug_(2,44) = 1.855, *p* = 0.168) (E) Manipulation of SST PL neurons do not affect sucrose binge drinking (F_drug_(2,46) = 0.162, *p* = 0.85, F_virus_(1,23) = 0.088, *p* = 0.768, F_virus x drug_(2,46) = 0.404, *p* = 0.669). (F) Schematic of electrophysiology recording in DREADD-expressing SST neurons and representative traces of RMP of hM3Dq- and KORD-expressing SST neurons during bath application of 10 uM CNO (top) and 1 uM SalB (bottom). (G) CNO depolarizes (baseline vs. washout, paired *t*(5) = 3.295, *p* = 0.022) and SalB hyperpolarizes (baseline vs. washout, paired *t*(6) = 4.473, *p* = 0.007) resting membrane potential of KORD- and hM3Dq-expressing SST neurons (CNO vs. SalB unpaired *t*(10) = 5.483, *p* < 0.001). (H) Schematic of electrophysiology recording in L2/3 pyramidal neurons and representative traces of sIPSC recordings in glutamate receptor blocker (3mM kynurenic acid)-containing aCSF before and after bath application of 10 μM CNO and 1 μM SalB. (I) CNO-induced activation of PL SST neurons increases sIPSC frequency on pyramidal neurons (baseline vs. washout, paired *t*(5) = 4.0, *p* = 0.016) with no changes in amplitude (paired *t*(5) = 1.12, *p* = 0.325). SalB-induced inhibition of PL SST neurons otherwise produced a decrease in sIPSC frequency (paired *t*(6) = 2.771, *p* = 0.039) and an increase in sIPSC amplitude (paired *t*(7) = 5.666, *p* = 0.002). * *p* < 0.05. ** *p* < 0.01, ***, *p* < 0.001, **** *p* < 0.0001. Tukey’s post-hoc test.

Surprisingly, both activation and inhibition of PL SST neurons (via CNO and SalB systemic administration, respectively) reduced alcohol consumption during the binge sessions, with no changes in alcohol drinking in viral control-injected mice (**Figure 2C**). This effect was seen in both sexes, although CNO-induced activation in females appeared to be less effective than SalB-induced inhibition (**Figure 2D**). Consumption of 10% sucrose DID (**Figure 2E**) was unaltered in DREADD-expressing and control mice, suggesting that the CNO- and SalB induced effect was specific to alcohol consumption in DREADD-expressing mice, and not generalizable to other rewarding fluids.

Next, we performed whole cell patch clamp on hM3Dq/KORD-expressing SST neurons, visualized by mCherry and mCitrine fluorescence respectively, to verify the functionality of the DREADDs. As expected, 10-min bath application of CNO produced a significant depolarization in the RPM of SST neurons, whereas SalB resulted in a significant hyperpolarization (**Figure 2F, G**). Voltage clamp experiments on layer 2/3 pyramidal neurons revealed an increase in sIPSC frequency following bath CNO application, with no change in amplitude (**Figure 2H, I**). Following bath application of SalB there was a decrease in sIPSC frequency, and interestingly, an increase in sIPSC amplitude (**Figure 2I**).

Overall, these data reveal a role of PL SST neurons in regulating binge-like alcohol intake by regulating the inhibitory dynamics onto pyramidal neurons, a major source of excitatory output in the PL.

### A sex-specific SST-mediated inhibitory circuit in the PL

Due to our behavioral effects showing that bidirectional chemogenetic manipulation of SST neurons resulted in similar reductions in alcohol consumption, we further investigated whether SST neurons and pyramidal neurons may interact both directly and indirectly via an intermediary GABAergic source, as suggested by Cummings and Clem (2019) (26). We injected SST-IRES-Cre male and female mice with the Cre-dependent viral construct encoding for the light-inducible channerhodopsin-2 (ChR2) in the PL (**Figure 3A**). Action potentials in ChR2+ SST neurons were elicited by 1-ms blue light (470 nm) photostimulation, with full action potential fidelity upward of 10 Hz (**Figure 3A**), consistent with SST firing patterns we have previously published (36). Next, we used a paired-pulse photostimulation protocol (2 x 1 ms pulses, 100 ms interval) to evoke release from SST neurons while recording from either pyramidal neurons or putatively GABAergic, non-SST neurons. The cell type identity was confirmed by membrane properties (capacitance and membrane resistance, **Supplementary Figure 3C, D**) and spiking characteristics (**Supplementary Figure 3E, F),** was consistent with the published literature (26). This stimulation protocol revealed direct, monosynaptic connections between SST neurons and both neighboring subpopulations, in which photostimulation of SST neurons induced potent, time-locked IPSCs in both pyramidal neurons and non-SST interneurons (**Figure 3B**).

**Figure 3:**
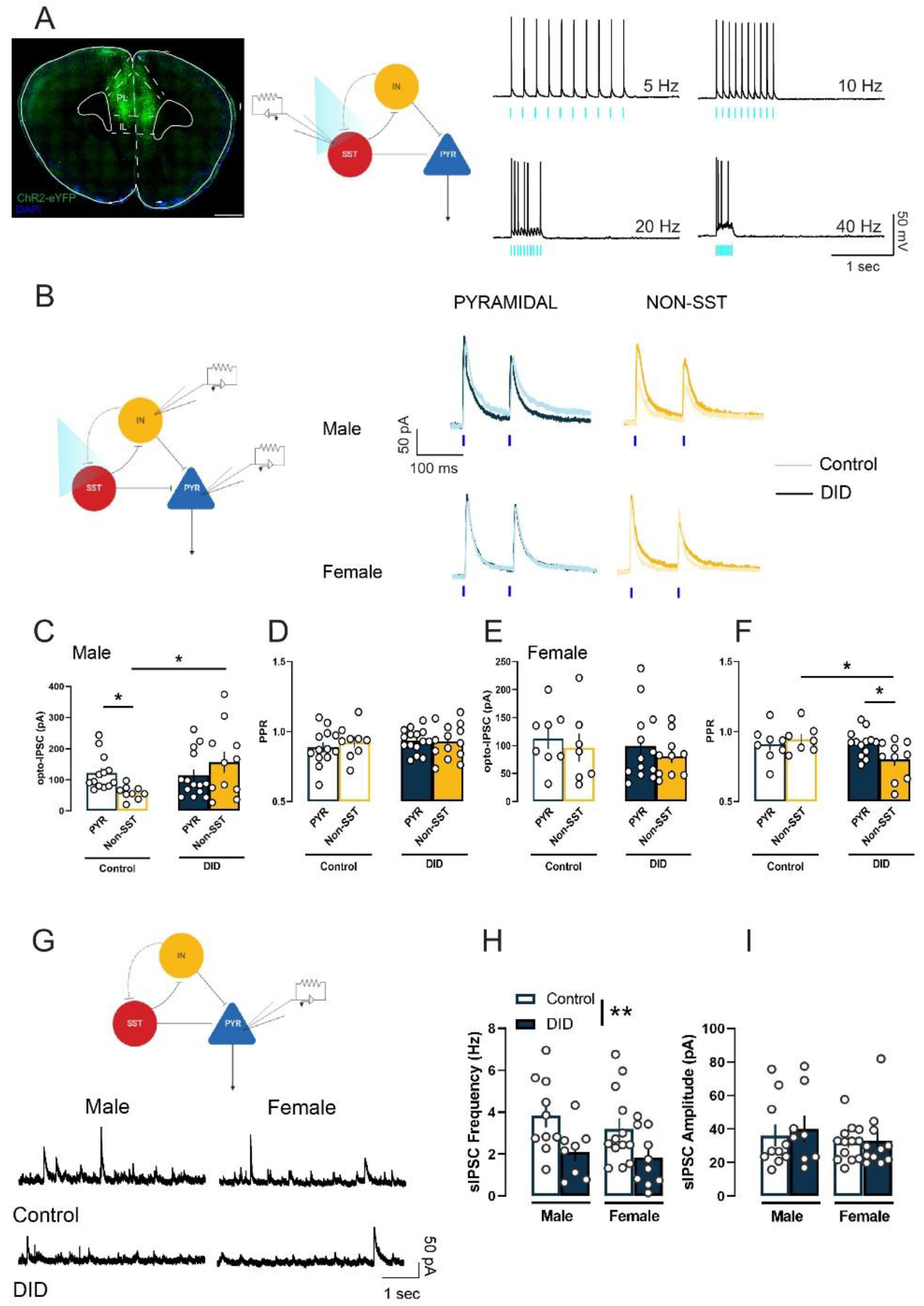
SST-mediated inhibitory dynamic in the PL is moderated by sex and alcohol binge drinking. (A) (Left) ChR2-expressing SST neurons in the PL. Scale bar 1mm. (Right) Photostimulated action potential fidelity in SST neurons upward to 10 Hz. (B) Representative traces of SST-mediated evoked IPSCs in PL pyramidal neurons and non-SST interneurons in males (top) and females (bottom). (C) In control males, SST cells provide stronger GABA_A_R-mediated inhibition onto pyramidal neurons than onto other non-SST, putatively GABAergic neurons (*p* = 0.016). Binge drinking reversed this dynamic, in which it increased SST-evoked IPSC amplitude onto non-SST neurons. No change in SST-evoked IPSC amplitude in pyramidal neurons following binge drinking (control non-SST v. DID non-SST: *p =* 0.011; control pyramidal: n = 13 cells, control non-SST: n = 9 cells, DID pyramidal: n = 14 cells, DID non-SST: n = 11 cells; F_cell type_(1,43) = 0.218, *p* = 0.642; F_binge_(1,43) = 4.109, *p* = 0.049, F_cell type x binge_(1,43) = 5.731, *p* = 0.021). (D) No differences in SST-evoked IPSC paired pulse ratio across cell types and drinking history in males (F_cell type_(1,43) = 0.271, *p* = 0.604; F_binge_(1,43) = 0.661, *p* = 0.420, F_cell type x binge_(1,43) = 0.407, *p* = 0.526). (E) In females, SST neurons provide balanced inhibition onto pyramidal neurons and non-SST neurons, and binge drinking did not affect this dynamic with no differences in SST-evoked IPSC peak amplitude (control pyramidal: n = 8 cells, control non-SST: n = 7 cells, DID pyramidal: n = 13 cells, DID non-SST: 10 cells; F_cell type_(1,34) = 0.849, *p* = 0.363; F_binge_(1,34) = 0.566, *p* = 0.457, F_cell type x binge_(1,34) = 0.002, *p* = 0.962). (F) Binge drinking decreased SST-evoked IPSC paired pulse ratio, a measure of neurotransmitter release probability, in female non-SST neurons (control non-SST vs. DID non-SST: *p* = 0.025; DID pyramidal vs. DID non-SST: *p* = 0.039; F_cell type_(1,34) = 1.11, *p* = 0.299; F_binge_(1,34) = 3.406, *p* = 0.073, F_cell type x binge_(1,34) = 4.13, *p* = 0.05). (G) Representative traces of sIPSC recordings in PL layer 2/3 pyramidal neurons. (H) Binge drinking reduces sIPSC frequency in male (DID: n = 8 cells, control: n = 10 cells) and female (DID: n = 10 cells, control: n = 13 cells) PL layer 2/3 pyramidal neurons (F_sex_(1,37) = 0.783, *p* = 0.381; F_binge_(1,37) = 9.558, *p* = 0.003, F_sex x binge_(1,37) = 0.132, *p* = 0.718). (I) sIPSC amplitude in male and female PL layer 2/3 pyramidal neurons was not altered following binge drinking (F_sex_(1,37) = 1.055, *p* = 0.31; F_binge_(1,37) = 0.199, *p* = 0.657, F_sex x binge_(1,37) = 0.065, *p* = 0.798). DID: N = 5 male mice and 5 female mice, control: N = 6 male mice and 4 female mice. * *p* < 0.05. ** *p* < 0.01. Bonferroni’s post-hoc test.

In control males, SST neuron-mediated inhibition is biased towards pyramidal neurons, with larger optically evoked IPSC in pyramidal neurons than in non-SST neurons (**Figure 3C**). However, alcohol binge drinking reversed this dynamic by increasing SST neuron-mediated IPSC amplitude onto non-SST neurons (**Figure 3C**). IPSC paired pulse ratio (PPR), a measurement of neurotransmitter release probability (37), are nevertheless similar across cell types in male PL circuit and not affected by alcohol binge drinking (**Figure 3D**).

In contrast to males, PL SST neurons in females showed a balanced inhibition dynamic between pyramidal neurons and non-SST neurons, in which optically evoked IPSC amplitudes are similar across cell types and drinking groups (**Figure 3E**). However, there is a decrease in SST neuron-mediated IPSC PPR onto non-SST neurons of DID female mice, as compared to control non-SST neurons and DID pyramidal neurons, suggesting an increase in GABA release probability from SST neurons onto other GABAergic populations (**Figure 3F**). In complementary experiments, we assessed spontaneous GABA transmission onto neighboring pyramidal cells. Both males and females had decreased sIPSC frequency on pyramidal neurons in layer 2/3 post-DID as compared to water controls (**Figure 3G, H**). sIPSC amplitudes were unaltered across groups (**Figure 3G, I**).

Together, these data provide evidence for a microcircuit of SST neurons in the PL to directly and indirectly gate the gain of pyramidal neurons to control binge-like alcohol drinking, with sex differences in the relative strength of SST neurons-mediated inhibition of excitatory pyramidal neurons and other GABAergic populations. Repeated cycles of excessive alcohol use disrupts these dynamics and disinhibits pyramidal neurons in the PL by enhancing SST-mediated inhibitory tone onto other GABAergic source, and sets the stage for the ability for diverse manipulations of SST neurons to produce similar behavioral changes.

### Direct inhibition of PL pyramidal neurons reduce binge-like alcohol drinking

Our results thus far point to a reciprocal relationship between inhibitory transmission in PL pyramidal neurons and binge-like alcohol intake. Next, we investigated whether direct silencing of PL pyramidal neurons is sufficient to curb alcohol drinking in the DID paradigm, consistent with our hypothesized overall circuit. We injected wildtype C57Bl6/J males and females in the PL with a viral construct encoding for the inhibitory Gi-coupled hM4D DREADD under the control of the CamKIIα promoter (representative viral injection and experimental timeline **Figure 4A**). PL pyramidal silencing decreased alcohol intake in both sexes (**Figure 4B**). There was no sex difference in responsiveness to pyramidal inhibition (**Figure 4C**). In sum, these data supports the overall framework of the PL as a critical hub for controlling aberrant alcohol consumption. Disruption of the excitation/inhibition dynamics within this microcircuitry can enhance susceptibility to binge-like alcohol drinking, and subsequent dependence and abuse.

**Figure 4:**
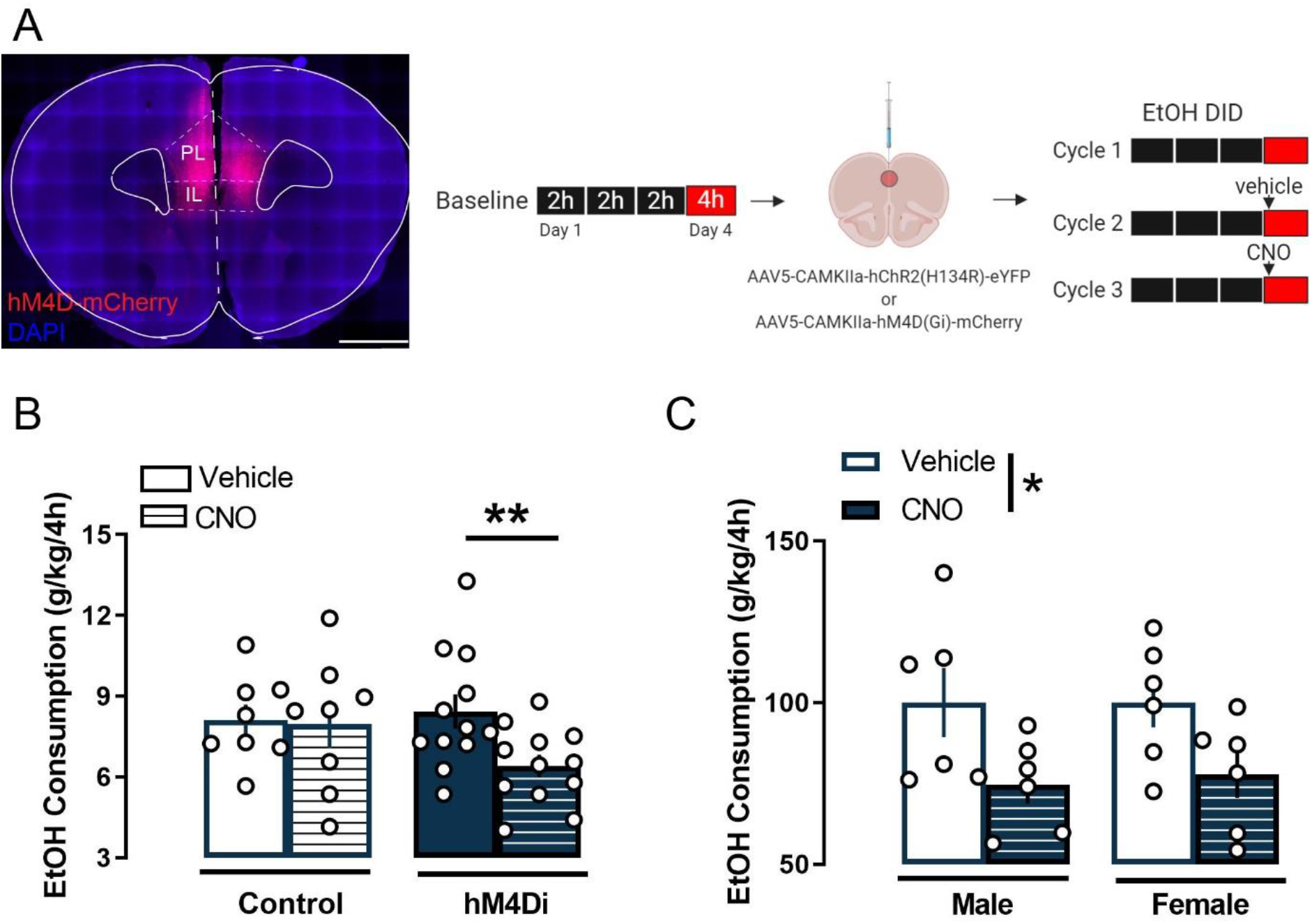
Direct silencing of PL pyramidal neurons reduces binge drinking. (A) (Left) Representative images of the hM4D(Gi) viral expression in PL pyramidal neurons. Scale bar 1mm. (Right) Schematic of the experimental design. Both male and female C57BL/6J mice were used in this experiment. (B) CNO-induced inhibition of PL pyramidal neurons decreases binge drinking in hM4D(Gi)-expressing mice, (vehicle vs. CNO, *p* = 0.005), but not in control virus mice (*p* > 0.99) (F_drug_(1,18) = 5.714, *p* = 0.028, F_virus_(1, 18) = 0.64, *p* = 0.434, F_virus x drug_(1,18) = 4.246, *p* = 0.050). (C) Inhibition of PL pyramidal neurons are equally effective in reducing binge drinking in both male and female mice (F_drug_(1,10) = 9.864, *p* = 0.01, F_sex_(1, 10) = 0.033, *p* = 0.858, F_sex x drug_(1,10) = 0.041, *p* = 0.843). * *p* < 0.05. ** *p* < 0.01. Bonferroni’s post-hoc test.

**Figure 5:**
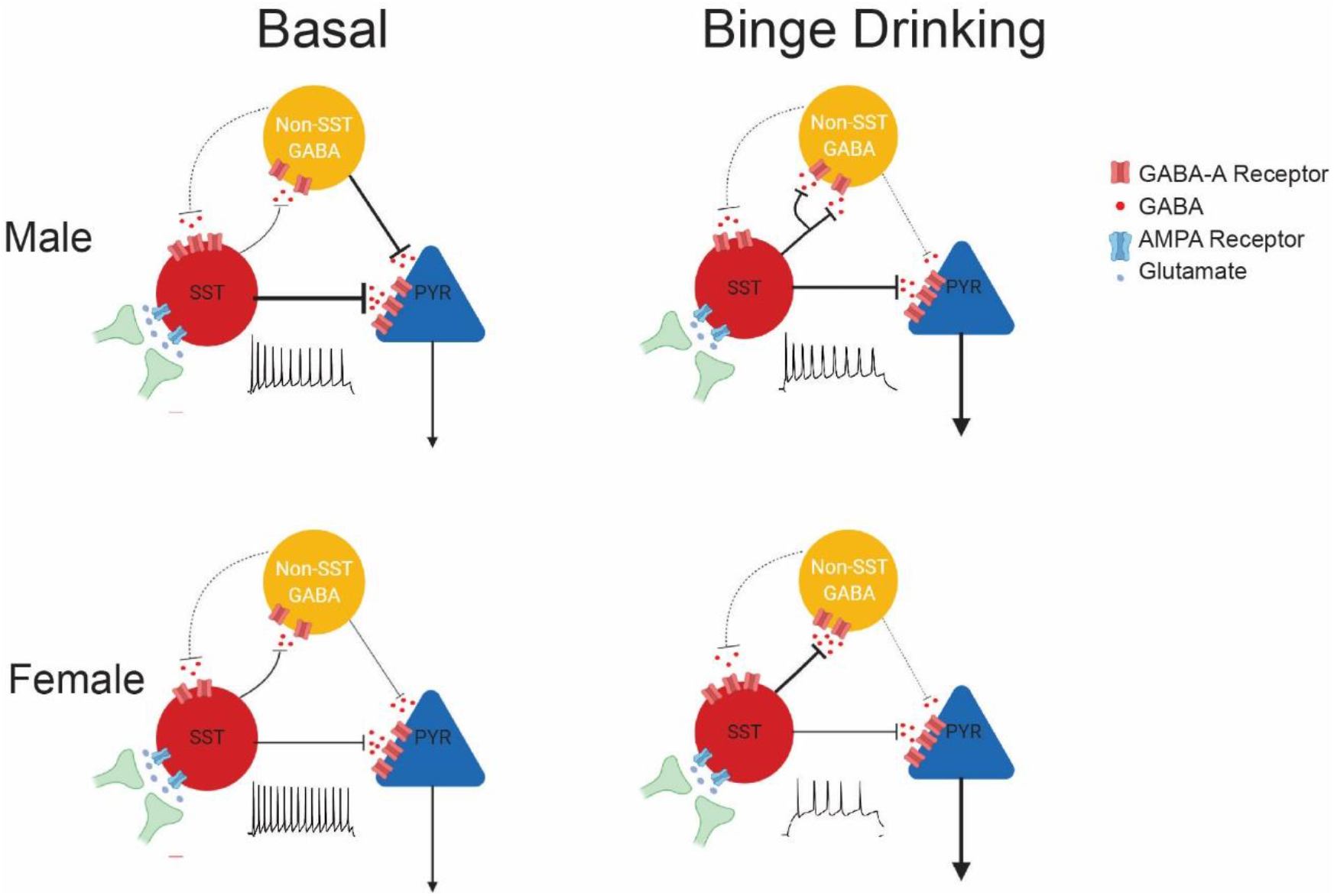
Microcircuitry of the PL cortex and binge drinking-induced plasticity. In basal states, males and females exhibit differences in spontaneous inhibitory transmission onto PL SST neurons, with higher sIPSC amplitude in males. Alcohol binge drinking disrupts the excitation/inhibition dynamic on SST neurons by dampening excitatory transmission and reversing the sex difference in inhibitory transmission. Concurrently, alcohol binge drinking dampens intrinsic excitability of PL SST neurons, resulting in a hypoactive state. PL SST neurons provide inhibition onto both pyramidal neurons and putative GABAergic neurons in the PL cortex, with bias towards pyramidal neurons in males only. Alcohol binge drinking reverses this dynamic by augmenting SST neuron inhibitory output onto other GABAergic population. Pyramidal inhibition is critical for curbing alcohol consumption behaviors, thus disinhibition of pyramidal could increase the risk of compulsive alcohol drinking and alcohol dependence.

## DISCUSSION

Here, we demonstrate evidence for the role of SST neurons in the PL cortex in binge-like alcohol consumption consumption. Our data expand upon previous reports on the involvement of this GABAergic population in a host of neuropsychiatric disorders, particularly depression (23,24) and anxiety (25). Repeated cycles of alcohol binge drinking altered intrinsic excitability and excitation/inhibition dynamics of PL SST neurons, resulting in a state of hypoactivity within this population (**Figure 1**). Bidirectional manipulation of PL SST neurons via co-expressed excitatory and inhibitory DREADDs paradoxically reduced alcohol consumption following both excitation and inhibition of these neurons, by acting directly via their monosynaptic connections on pyramidal neurons and indirectly via neighboring GABAergic populations respectively (**Figures 2, 3**). The SST neurons-mediated inhibition dynamic in the PL is sex-dependent, favoring pyramidal neurons over putatively GABAergic, non-SST neurons in males and not females. Alcohol binge drinking potentially disinhibits pyramidal neurons by augmenting inhibitory strength of SST neurons onto other non-SST neurons (**Figure 3**). Lastly, facilitating pyramidal inhibition could mimic the effect of SST neuron manipulation in reducing binge drinking, providing a secondary approach for reducing alcohol consumption via this circuit (**Figure 4**). Overall, these results confirm and expand upon the role of the PL cortex as a critical neural hub for the development of excessive alcohol consumption, and SST neurons as a key regulator of local PL network activity.

Previous reports have demonstrated the multitude of neural adaptations in the PFC following alcohol dependence and withdrawal. In separate complementary studies, male C57BL6/J mice exposed to the chronic intermittent ethanol vapor (CIE) model of alcohol dependence exhibited elevated intrinsic excitability in layer 2/3 pyramidal neurons (21,22), increased dendritic spine density (22,38), and upregulated NMDA receptor expression (38). The DID paradigm, in contrast to the CIE model, models a binge drinking pattern that produces a highly intoxicated state without inducing dependence symptomologies (28), and changes seen in this model could be considered precursors to those that occurring following CIE. Previously, we demonstrated that four cycles of DID reduced glutamatergic transmission onto female pyramidal neurons in the PL via a reduction in cell-surface AMPA and NMDA receptors (20), which is not evident in males. Here we observed that the effects of DID on neuronal intrinsic excitability and synaptic transmission in the PL extends to SST neurons, in which SST neurons displayed a state of hypoactivity with diminished intrinsic excitability and a disrupted excitatory/inhibitory transmission dynamic in both sexes. Similar effects were observed in an intermittent-access model of alcohol drinking (39). Of note, despite a reduction in sEPSC frequency on SST neurons in both sexes, there is an increase in evoked GABA release probability, as evidenced by reduced IPSC paired-pulse ratio (39), suggesting that there could be a compensatory mechanism on a whole-circuit level to counteract the effects on excitatory transmission in single neurons. Alterations in sEPSC frequency indicate the effects of DID on a presynaptic locus, which could arise from a wide array of glutamatergic sources such as the basolateral amygdala (BLA), the ventral hippocampus, the thalamus as well as within the medial prefrontal cortex itself (12,40,41). Inputs from the BLA, a key regulator of behavioral stress response, to PL pyramidal neurons have been shown to be sufficient to drive anxiety-like behaviors in male C57BL/6J mice, and chronic stress can potentiate BLA-to-PFC synapses (11,42). The infralimbic cortex (IL) and the thalamus have been implicated in extinction of drug-seeking behavior (43) and response to alcohol cue (41), respectively. Further investigation into the effects of DID on excitatory transmission onto PL SST neurons at a synapse-specific level is needed to fully reveal how alcohol binge drinking-mediated dysregulation of glutamatergic inputs in PL SST neurons drives withdrawal-related negative affect related behaviors and escalation in drinking.

The altered excitatory synaptic inputs were accompanied by a sex-dependently opposing effects of alcohol binge drinking on inhibitory transmission onto PL SST neurons. We observed a basal sex difference in phasic sIPSC amplitude in PL SST neurons of non-drinking mice. Following DID, sIPSC amplitude was enhanced in female yet reduced in male SST neurons. These data overall suggest a substantial variability in the GABA_A_ receptor expression in these neurons, which interacts with alcohol binge drinking in a sex-dependent manner.

We previously reported that PL SST neurons are hyperexcitable following prolonged withdrawal from alcohol drinking (> 4 weeks) in a 2-bottle choice model (34), which contrasts with the hypoexcitable state in acute withdrawal (24-h) from DID observed here. It should be noted that the excitability recording in the former report was done following an acute stressor (forced swim). Other stress paradigms, such as fear conditioning (26) and morphine withdrawal (44),, could augment excitatory drive in PL SST neurons. These findings together suggest an extensive capacity for plasticity in these neurons that could be moderated by alcohol withdrawal stage and stress, a major predictor for relapse in alcohol drinking. The cellular and molecular mechanism underlying these alterations in SST intrinsic excitability are still unclear. Ethanol exposure has been shown to modify expression of key modulators of neuronal excitability, including ionotropic glutamate receptors (20,38), inward-rectifying potassium channel (45), voltage-gated calcium channels (46), and hyperpolarization-activated cation channels (HCN; (47). Intermittent access to alcohol increased medium afterhyperpolarization in PL SST neurons (39), which might be mediated by M-type potassium channels and HCN channels (48). Notably, in females following DID, PL SST neurons displayed highly depolarized resting membrane potential yet could not efficiently produce action potentials, suggesting a state of depolarization block. One hypothesis is that DID disrupts potassium exchange within these neurons via downregulation of potassium channel function, which has been observed in other GABAergic population (45) and leads to deficits in repolarization and action potential firings. These phenomena merit further investigation to fully demonstrate the alcohol binge drinking-induced cellular and molecular adaptations.

A major and unexpected finding in our study is that bidirectional manipulation of SST neurons activity via both an excitatory and inhibitory DREADDs similarly reduced alcohol binge drinking. CNO-mediated activation of SST neurons increased sIPSC frequency onto pyramidal neurons, concordant with a direct, monosynaptic SST to pyramidal neuron connection. On the other hand, SalB-induced silencing of SST neurons both decreased sIPSC frequency, while also increasing sIPSC amplitude onto pyramidal neurons. Our ChR2-assisted circuit mapping data revealed that SST neurons in the PL synapse extensively to neighboring populations, including pyramidal neurons and other GABA neurons. DREADD-induced silencing of SST neurons may lead to disinhibition of another local GABAergic source to maintain overall pyramidal inhibition, as evidenced by the increase in sIPSC amplitude in pyramidal neurons, although we cannot fully rule out an uncharacterized pharmacological action of Salvinorin B in the PL circuit. The intermediate GABAergic population is likely parvalbumin-expressing neurons, which comprise the majority of cortical interneurons and provide powerful inhibition of pyramidal output in the PL (26). A similar microcircuitry motif has been observed in the somatosensory cortex, in which photoinhibition of SST neurons decreases pyramidal firings in layer 4 (49). In males, barrel cortical SST neurons bias towards fast-spiking interneurons over pyramidal neurons. Xu et al. also identified a layer-specific SST-mediated circuitry using dual patch clamp, in which layer 2/3 SST neurons otherwise favor pyramidal inhibition (49). It is possible that our whole-field photostimulation approach targeted most GABAergic neuron-biased SST synapses in layer 2/3 of the PL. Additionally, we further showed that this PL microcircuit might be sex-dependent, in which SST neurons-mediated biased inhibition of pyramidal neurons over other putatively GABAergic neurons were observed in males only. This suggests that in females, SST neurons-mediated inhibition dynamic in the PL is balanced across different cell types. Whether the sex differences in inhibitory dynamic in the PL could partially contribute to the higher drinking levels and higher susceptibility to alcohol-related negative affective states in females merit further examination.

Additionally, we observed that DID reversed this inhibition dynamic and disinhibits pyramidal neurons in the PL by augmenting SST neuron-mediated inhibition onto other putatively GABAergic populations. Similar disinhibitory effects have been observed following fear conditioning (26), social fear (27) and morphine withdrawal (44), in which SST neurons-mediated IPSC amplitude onto parvalbumin neurons is enhanced with a reduction in IPSC PPR. This effect is accompanied by either an increase in action potential firing or excitatory inputs in PL SST neurons (26,44). Interestingly, our optogenetic-assisted circuit mapping results, combined with our data on attenuated action potential firing and sEPSC frequency in PL SST neurons, suggested that alcohol may produce complex and changes in the PL cortex that must be further teased apart. One interpretation is that the population-wide, optically-evoked IPSC transmission from SST neurons (**Figure 3**) does not reflect the changes on a single-neuron level in the intrinsic excitability experiments (**Figure 1**). Population-wide increase in SST-mediated inhibitory outputs may occur to compensate the reduction in excitation of single neurons, which bears resemblance to the discrepancy between spontaneous and evoked EPSC on SST neurons in the intermittent-access model (39). In addition, assessments of raw optogenetically evoked IPSC amplitudes must be interpreted with caution, as mouse-to-mouse variability in ChR2 injection may drive changes in overall results.

Other models of alcohol exposure have corroborated the effects of alcohol on excitatory and inhibitory inputs in PL pyramidal neurons. Intragastric administration of high ethanol doses in rats (5 g/kg) diminished GABA_A_ receptor-mediated inhibitory currents in PFC layer 5/6 pyramidal neurons accompanied by a reduction in expression of the α1 GABA_A_ receptor subunit (50). As pyramidal neurons are the main group of projection neurons from the PL, providing strong glutamatergic input to associated cortical regions (6,7), nucleus accumbens (10), ventral tegmental area (51) and amygdala (52), which are critically involved in emotional and motivated behaviors. Binge drinking-induced aberrant inhibitory inputs onto pyramidal neurons thus may dysregulate cortical control of downstream targets, enhancing susceptibility to and development of substance dependence and abuse. A recent report by Siciliano et al. (2019) found a prefrontal cortex (PFC)-to-periaquaductal gray circuit that controls compulsive binge drinking, and demonstrated that inhibitory activity of this PFC subpopulation during pre-dependent stage predicts future vulnerability to compulsive-like, punishment-resistant alcohol seeking behaviors (53). Here we found that direct silencing of pyramidal neurons via hM4Di was sufficient to curb alcohol binge drinking, supporting the overall framework of PL excitatory output driving alcohol consumption.

While our data demonstrate the extensive interaction between PL SST neurons and alcohol binge drinking, these neurons are not a unique modulator of excessive alcohol consumption. The cortex includes a diverse array of GABAergic neurons with distinct neurochemical and electrophysiological profiles, including NPY, DYN, and CRF (49), and their roles in neuropsychiatric disease are well-characterized. Additionally, the current study have yet to address the involvement of neuropeptide signaling, especially that of somatostatin, in modulating alcohol consumption. Previous reports have demonstrated that DID reduces NPY expression in the PL, and conversely intra-PL administration of NPY1 agonist and CRF1 antagonist could reduce alcohol binge drinking (17,16). Here we did not assess whether the effects of hM3Dq-induced excitation of PL SST neurons on alcohol binge drinking could be partially attributed to SST peptide release and SST signaling in the PL. The neuropeptide SST has been shown to act largely via a family of Gi/o-coupled receptors (SSTR1-5) to inhibit neuronal excitability (18,19). Future studies should examine the effects of alcohol binge drinking on somatostatin signaling in the PL, and the potential of somatostatin peptide as a therapeutic candidate for excessive alcohol consumption.

Overall, these results posit alterations in prelimbic SST neurons as a potential neural substrate of binge drinking. These neurons emerge as a strong candidate for treatment of AUD as they are well-situated to exert inhibition over the local PL circuitry, thereby controlling the flow of input and output signals of this region.

## ACKNOWLEDGEMENTS

The experiments were funded by The Brain and Behavior Research Foundation (NARSAD Young Investigator Award to NAC), NIAAA (1R21AA028088-01 to NAC) and Penn State’s Social Science Research Institute (Level 2 Award to NAC). NCD and NAC conceived of and designed the experiments, NCD, DFB, and MSN conducted the experiments, NCD and NAC analyzed and interpreted the experiments, and NCD and NAC wrote the paper with feedback from all authors. The authors would like to thank Dr. Janine Kwapis (Department of Biology, Pennsylvania State University) for her assistance with imaging, and Dr. Max Joffe for his critical comments on the manuscript.

## DISCLOSURES

The authors have no financial disclosures and no conflicts of interest to report.

---

**Supplementary Figure 1.**
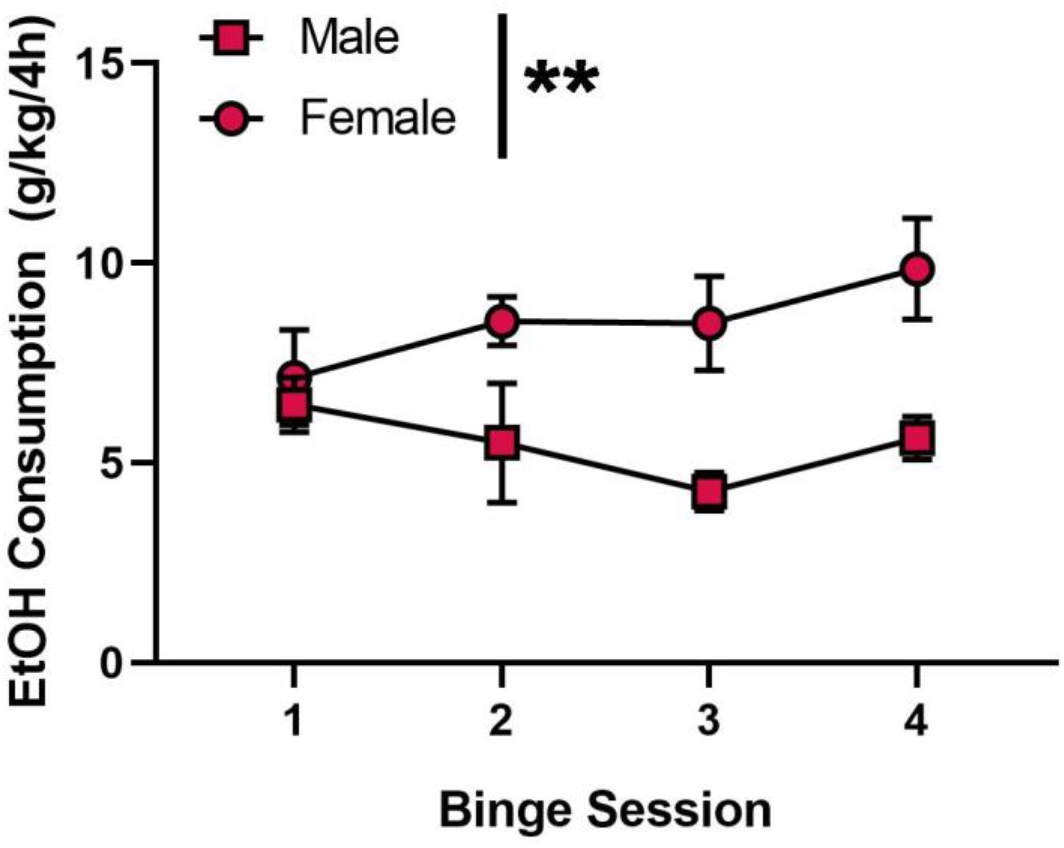
Female mice engage in higher level of alcohol binge drinking than male mice, related to Figure 1. Average consumption of 20% (v/v) ethanol solution in 4-h binge session in Drinking-in-the-Dark (DID) paradigm showed a sex difference (N = 6 males and 5 females, F_sex_(1,9) = 15.07, *p* = 0.003), and these levels are stable across cycles (F_cycle_(3,27) = 0.713, *p* = 0.552, F_sex x binge_(3,27) = 1.561, *p* = 0.221)

**Supplementary Figure 2.**
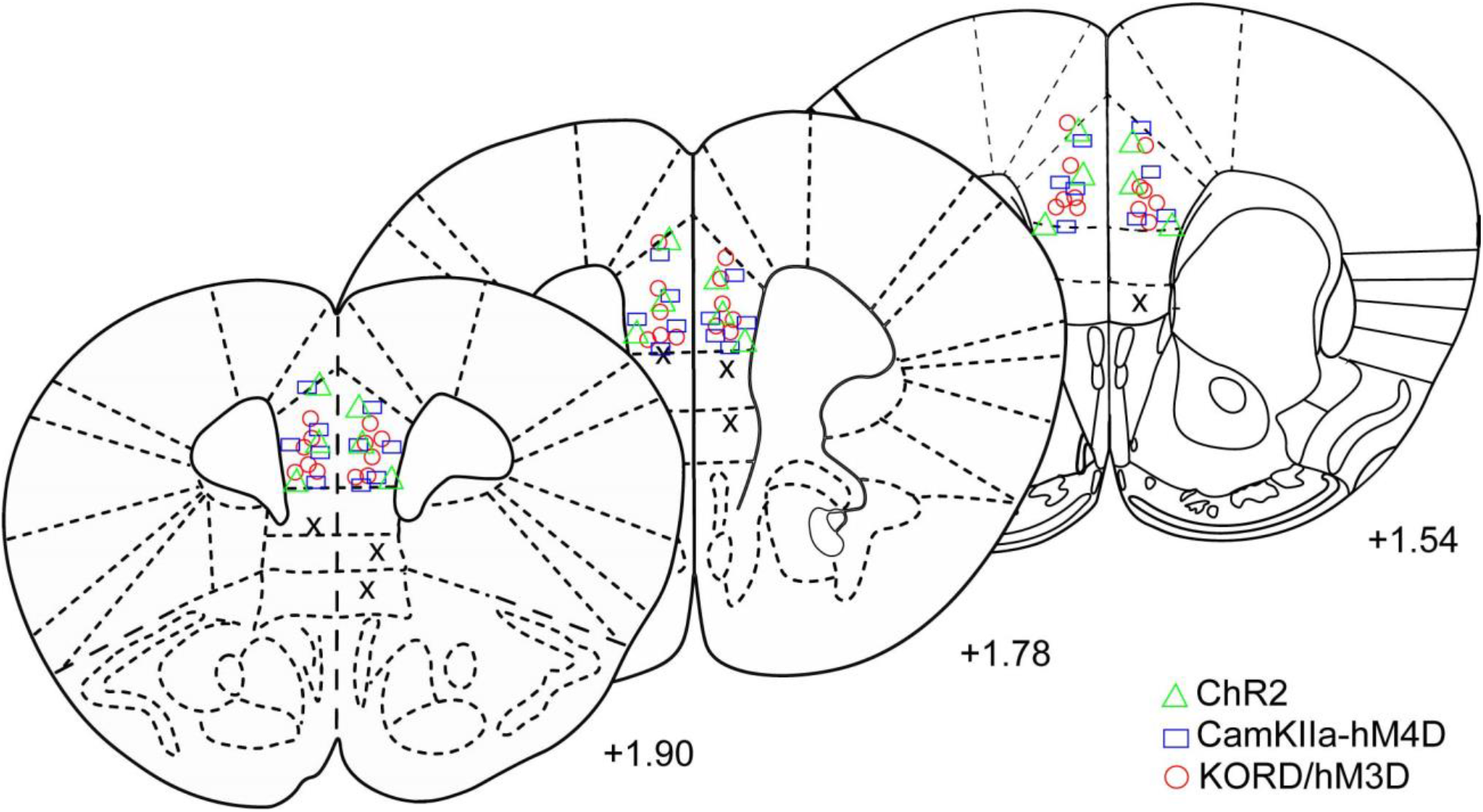
Histological verification of viral expression in the PL, related to figures 2, 3 and 4. Each symbol represents the center of injections, where the expression of fluorophore-tagged virus is the brightest. Red circles: KORD/hM3Dq mice. Green triangles: ChR2 mice. Blue rectangles: CamKIIa promoter-hM4D mice. X: missed injection.

**Supplementary Figure 3.**
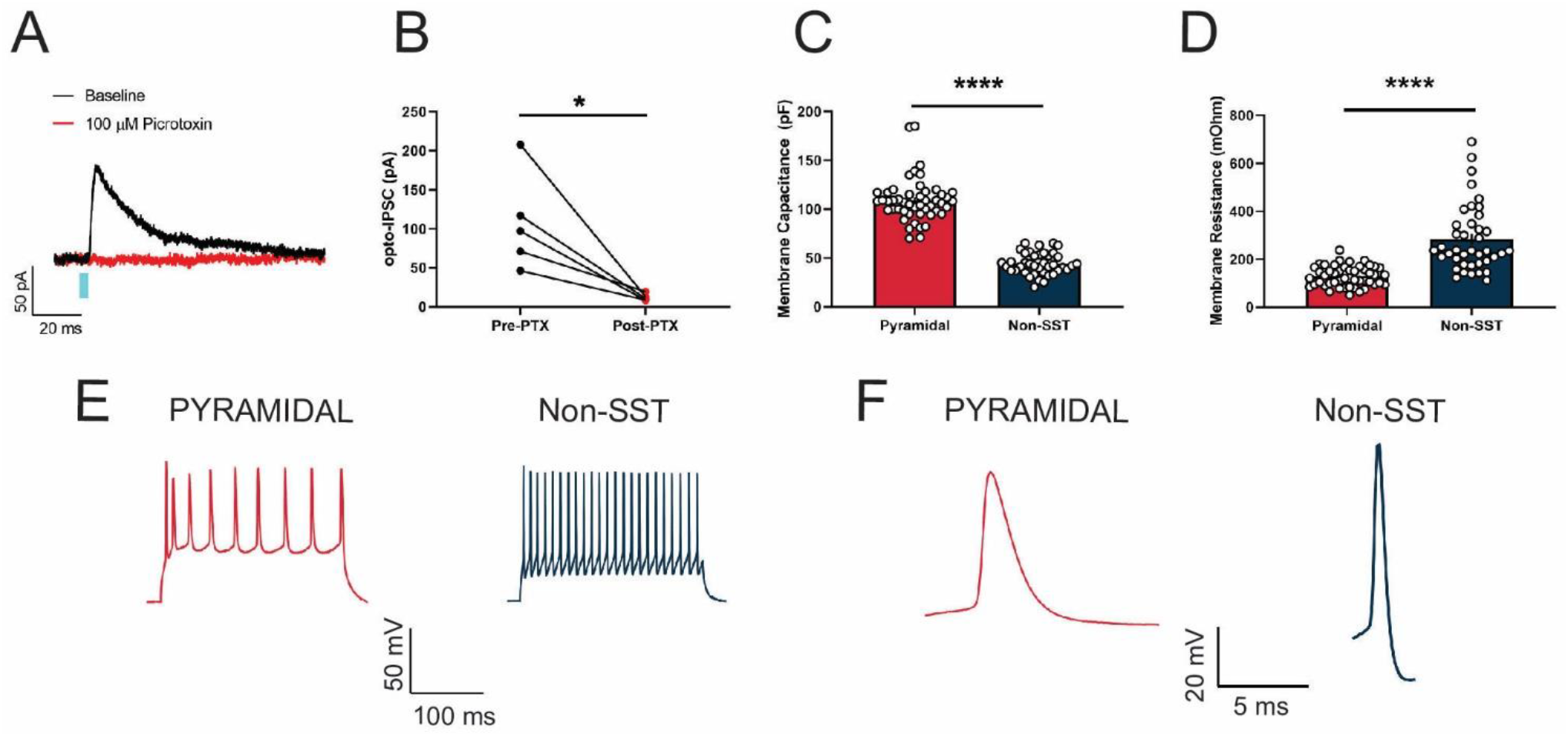
Membrane properties of pyramidal neurons and non-SST neurons in the PL, related to Figure 3. (A) Representative traces of outward currents in pyramidal neurons in the PL following photostimulation of SST neurons during baseline (black) and after bath application of the GABA_A_ receptor antagonist picrotoxin. (B) Bath application of 100 μM picrotoxin completely abolished SST-evoked IPSC in pyramidal neurons (paired *t*(4) = 3.465, *p* = 0.025). (C) Pyramidal neurons in the PL exhibit distinctively higher membrane capacitance than non-SST neurons (unpaired *t*(87) = 16.84, *p* < 0.0001). (D) Pyramidal neurons in the PL exhibit lower membrane resistance than non-SST neurons (unpaired *t*(87) = 7.396, *p* < 0.0001). (E) Representative traces of action potentials in pyramidal neurons and non-SST neurons in response to a +200 pA somatic current injection at RMP. (F) Representative single action potentials in pyramidal neurons and non-SST neurons. Cells were classified as pyramidal neurons with long action potential width (>6 ms), or as non-SST neurons with short action potential width (1-3 ms).

## Notes

### Competing Interest Statement

The authors have declared no competing interest.

## References

1. National Institute of Alcohol Abuse and Alcoholism. (2020). Drinking Levels Defined. National Institute of Alcohol Abuse and Alcoholism. https://www.niaaa.nih.gov/alcohol-health/overview-alcohol-consumption/moderate-binge-drinking

2. American Psychiatric Association. (2013). Diagnostic and Statistical Manual of Mental Disorders, 5th. Ed. American Psychiatric Publishing.

3. National Institute of Alcohol Abuse and Alcoholism. (2020). Alcohol Facts and Statistics. NIAAA [WWW Document]. https://www.niaaa.nih.gov/publications/brochures-and-fact-sheets/alcohol-facts-and-statistics

4. Wilsnack, R. W., Wilsnack, S. C., Gmel, G., & Kantor, L. W. (2017). Gender differences in binge drinking prevalence, predictors, and consequences. Alcohol Research: Current Reviews, 39(1), 57–76.

5. Abrahao, K. P., Salinas, A. G., & Lovinger, D. M. (2017). Alcohol and the Brain: Neuronal Molecular Targets, Synapses, and Circuits. Neuron, 96(6), 1223–1238. https://doi.org/10.1016/j.neuron.2017.10.032

6. Koob, G. F., & Volkow, N. D. (2016). Neurobiology of addiction: a neurocircuitry analysis. The Lancet Psychiatry, 3(8), 760–773. https://doi.org/10.1016/S2215-0366(16)00104-8

7. George, O., & Koob, G. F. (2010). Individual differences in prefrontal cortex function and the transition from drug use to drug dependence. Neuroscience and Biobehavioral Reviews, 35(2), 232–247. https://doi.org/10.1016/j.neubiorev.2010.05.002

8. Crowley, Nicole A., Bloodgood, D. W., Hardaway, J. A., Kendra, A. M., McCall, J. G., Al-Hasani, R., McCall, N. M., Yu, W., Schools, Z. L., Krashes, M. J., Lowell, B. B., Whistler, J. L., Bruchas, M. R., & Kash, T. L. (2016). Dynorphin controls the gain of an amygdalar anxiety circuit. Cell Reports, 14(12), 2774–2783. https://doi.org/10.1016/j.celrep.2016.02.069

9. Hwa, L. S., Neira, S., Flanigan, M. E., Stanhope, C. M., Pina, M. M., Pati, D., Hon, O. J., Yu, W., Kokush, E., Calloway, R., Boyt, K., & Kash, T. L. (2020). Alcohol drinking alters stress response to predator odor via bnst kappa opioid receptor signaling in male mice. ELife, 9, e59709. https://doi.org/10.7554/eLife.59709

10. Britt, J. P., Benaliouad, F., McDevitt, R. A., Stuber, G. D., Wise, R. A., & Bonci, A. (2012). Synaptic and behavioral profile of multiple glutamatergic inputs to the nucleus cccumbens. Neuron, 76(4), 790–803. https://doi.org/10.1016/j.neuron.2012.09.040

11. Lowery-Gionta, E. G., Crowley, N. A., Bukalo, O., Silverstein, S., Holmes, A., & Kash, T. L. (2018). Chronic stress dysregulates amygdalar output to the prefrontal cortex. Neuropharmacology, 139, 68–75. https://doi.org/10.1016/j.neuropharm.2018.06.032

12. McGarry, L. M., & Carter, A. G. (2016). Inhibitory gating of basolateral Amygdala inputs to the prefrontal cortex. Journal of Neuroscience, 36(36), 9391–9406. https://doi.org/10.1523/JNEUROSCI.0874-16.2016

13. Jin, J., & Maren, S. (2015m). Prefrontal-hippocampal interactions in memory and emotion. Frontiers in Systems Neuroscience, 9(170). https://doi.org/10.3389/fnsys.2015.00170

14. Tejeda, H. A., Counotte, D. S., Oh, E., Ramamoorthy, S., Schultz-Kuszak, K. N., Bäckman, C. M., Chefer, V., O’Donnell, P., & Shippenberg, T. S. (2013). Prefrontal cortical kappa-opioid receptor modulation of local neurotransmission and conditioned place aversion. Neuropsychopharmacology, 38(9), 1770–1779. https://doi.org/10.1038/npp.2013.76

15. George, O., Sanders, C., Freiling, J., Grigoryan, E., Vu, S., Allen, C. D., Crawford, E., Mandyam, C. D., & Koob, G. F. (2012). Recruitment of medial prefrontal cortex neurons during alcohol withdrawal predicts cognitive impairment and excessive alcohol drinking. Proceedings of the National Academy of Sciences of the United States of America, 109(44), 18156–18161. https://doi.org/10.1073/pnas.1116523109

16. Robinson, S. L., Perez-Heydrich, C. A., & Thiele, T. E. (2019). Corticotropin releasing factor type 1 and 2 receptor signaling in the medial prefrontal cortex modulates Binge-like ethanol consumption in C57BL/6J mice. Brain Sciences, 9(7), 171. https://doi.org/10.3390/brainsci9070171

17. Robinson, S. L., Marrero, I. M., Perez-Heydrich, C. A., Sepulveda-Orengo, M. T., Reissner, K. J., & Thiele, T. E. (2019). Medial prefrontal cortex neuropeptide Y modulates binge-like ethanol consumption in C57BL/6J mice. Neuropsychopharmacology, 44(6), 1132–1140. https://doi.org/10.1038/s41386-018-0310-7

18. Urban-Ciecko, J., & Barth, A. L. (2016). Somatostatin-expressing neurons in cortical networks. Nature Reviews Neuroscience, 17(7), 401–409. https://doi.org/10.1038/nrn.2016.53

19. Brockway, D. F., & Crowley, N. A. (2020). Turning the ‘Tides on neuropsychiatric diseases: The role of peptides in the prefrontal cortex. Frontiers in Behavioral Neuroscience, 14. https://doi.org/10.3389/fnbeh.2020.588400

20. Crowley, N.A., Magee, S. N., Feng, M., Jefferson, S. J., Morris, C. J., Dao, N. C., Brockway, D. F., & Luscher, B. (2019). Ketamine normalizes binge drinking-induced defects in glutamatergic synaptic transmission and ethanol drinking behavior in female but not male mice. Neuropharmacology, 149, 35–44. https://doi.org/10.1016/j.neuropharm.2019.02.003

21. Pleil, K. E., Lowery-Gionta, E. G., Crowley, N. A., Li, C., Marcinkiewcz, C. A., Rose, J. H., McCall, N. M., Maldonado-Devincci, A. M., Morrow, A. L., Jones, S. R., & Kash, T. L. (2015). Effects of chronic ethanol exposure on neuronal function in the prefrontal cortex and extended amygdala. Neuropharmacology, 99, 735–749. https://doi.org/10.1016/j.neuropharm.2015.06.017

22. Varodayan, F. P., Sidhu, H., Kreifeldt, M., Roberto, M., & Contet, C. (2018). Morphological and functional evidence of increased excitatory signaling in the prelimbic cortex during ethanol withdrawal. Neuropharmacology, 133, 470–480. https://doi.org/10.1016/j.neuropharm.2018.02.014

23. Fee, C., Banasr, M., & Sibille, E. (2017). Somatostatin-positive gamma-aminobutyric acid interneuron deficits in depression: cortical microcircuit and therapeutic perspectives. Biological Psychiatry, 82(8), 549–559. https://doi.org/10.1016/j.biopsych.2017.05.024

24. Fuchs, T., Jefferson, S. J., Hooper, A., Yee, P. H., Maguire, J., & Luscher, B. (2017). Disinhibition of somatostatin-positive GABAergic interneurons results in an anxiolytic and antidepressant-like brain state. Molecular Psychiatry, 22(6), 920–930. https://doi.org/10.1038/mp.2016.188

25. Soumier, A., & Sibille, E. (2014). Opposing effects of acute versus chronic blockade of frontal cortex somatostatin-positive inhibitory neurons on behavioral emotionality in mice. Neuropsychopharmacology, 39(9), 2252–2262. https://doi.org/10.1038/npp.2014.76

26. Cummings, K. A., & Clem, R. L. (2020). Prefrontal somatostatin interneurons encode fear memory. Nature Neuroscience, 23(1), 61–74. https://doi.org/10.1038/s41593-019-0552-7

27. Xu, Haifeng, Liu, L., Tian, Y., Wang, J., Li, J., Zheng, J., Zhao, H., He, M., Xu, T. Le, Duan, S., & Xu, H. (2019). A disinhibitory microcircuit mediates conditioned social fear in the prefrontal cortex. Neuron, 102(3), 668–682. https://doi.org/10.1016/j.neuron.2019.02.026

28. Crowley, Nicole A., Dao, N. C., Magee, S. N., Bourcier, A. J., & Lowery-Gionta, E. G. (2019). Animal models of alcohol use disorder and the brain: From casual drinking to dependence. Translational Issues in Psychological Science, 5(3), 222–242. https://doi.org/10.1037/tps0000198

29. Rhodes, J. S., Best, K., Belknap, J. K., Finn, D. A., & Crabbe, J. C. (2005). Evaluation of a simple model of ethanol drinking to intoxication in C57BL/6J mice. Physiology and Behavior, 84(1), 53–63. https://doi.org/10.1016/j.physbeh.2004.10.007

30. Vardy, E., Robinson, J. E., Li, C., Olsen, R. H. J., DiBerto, J. F., Giguere, P. M., Sassano, F. M., Huang, X. P., Zhu, H., Urban, D. J., White, K. L., Rittiner, J. E., Crowley, N. A., Pleil, K. E., Mazzone, C. M., Mosier, P. D., Song, J., Kash, T. L., Malanga, C. J., … Roth, B. L. (2015). A new DREADD facilitates the multiplexed chemogenetic interrogation of behavior. Neuron, 86(4), 936–946. https://doi.org/10.1016/j.neuron.2015.03.065

31. Li, H., Penzo, M. A., Taniguchi, H., Kopec, C. D., Huang, Z. J., & Li, B. (2013). Experience-dependent modification of a central amygdala fear circuit. Nature Neuroscience, 16(3), 332–339. https://doi.org/10.1038/nn.3322

32. Lee, J. H., Durand, R., Gradinaru, V., Zhang, F., Goshen, I., Kim, D. S., Fenno, L. E., Ramakrishnan, C., & Deisseroth, K. (2010). Global and local fMRI signals driven by neurons defined optogenetically by type and wiring. Nature, 465(7299), 788–792. https://doi.org/10.1038/nature09108

33. Rinker, J. A., Marshall, S. A., Mazzone, C. M., Lowery-Gionta, E. G., Gulati, V., Pleil, K. E., Kash, T. L., Navarro, M., & Thiele, T. E. (2017). Extended amygdala to ventral tegmental area corticotropin-releasing factor circuit controls binge ethanol intake. Biological Psychiatry, 81(11), 930–940. https://doi.org/10.1016/j.biopsych.2016.02.029

34. Dao, N. C., Suresh Nair, M., Magee, S. N., Moyer, J. B., Sendao, V., Brockway, D. F., & Crowley, N. A. (2020). Forced abstinence from alcohol induces sex-specific depression-like behavioral and neural adaptations in somatostatin neurons in cortical and amygdalar regions. Frontiers in Behavioral Neuroscience, 14(86). https://doi.org/10.3389/fnbeh.2020.00086

35. Ting, J. T., Lee, B. R., Chong, P., Soler-Llavina, G., Cobbs, C., Koch, C., Zeng, H., & Lein, E. (2018). Preparation of Acute Brain Slices Using an Optimized N-Methyl-D-glucamine Protective Recovery Method. Journal of Visualized Experiments : JoVE, 132, 53825. https://doi.org/10.3791/53825

36. Dao, N. C., Brockway, D. F., & Crowley, N. A. (2019). In vitro optogenetic characterization of neuropeptide release from prefrontal cortical somatostatin neurons. Neuroscience, 419, 1–4. https://doi.org/10.1016/j.neuroscience.2019.08.014

37. Choi, S., & Lovinger, D. M. (1997). Decreased probability of neurotransmitter release underlies striatal long-term depression and postnatal development of corticostriatal synapses. Proceedings of the National Academy of Sciences of the United States of America, 94(6), 2665–2670. https://doi.org/10.1073/pnas.94.6.2665

38. Kroener, S., Mulholland, P. J., New, N. N., Gass, J. T., Becker, H. C., & Chandler, L. J. (2012). Chronic alcohol exposure alters behavioral and synaptic plasticity of the rodent prefrontal cortex. PLoS ONE, 7(5). https://doi.org/10.1371/journal.pone.0037541

39. Joffe, M. E., Winder, D. G., & Conn, P. J. (2020). Contrasting sex-dependent adaptations to synaptic physiology and membrane properties of prefrontal cortex interneuron subtypes in a mouse model of binge drinking. Neuropharmacology, 178, 108126. https://doi.org/10.1016/j.neuropharm.2020.108126

40. Penzo, M. A., Robert, V., & Li, B. (2014). Fear conditioning potentiates synaptic transmission onto long-range projection neurons in the lateral subdivision of central amygdala. Journal of Neuroscience, 34(7), 2432–2437. https://doi.org/10.1523/JNEUROSCI.4166-13.2014

41. Fuchs, R. A., Evans, K. A., Ledford, C. C., Parker, M. P., Case, J. M., Mehta, R. H., & See, R. E. (2005). The role of the dorsomedial prefrontal cortex, basolateral amygdala, and dorsal hippocampus in contextual reinstatement of cocaine seeking in rats. Neuropsychopharmacology, 30(2), 296–309. https://doi.org/10.1038/sj.npp.1300579

42. Marcus, D. J., Bedse, G., Gaulden, A., Ryan, J. D., Clauss, J., Delpire, E., Lee, F. S., Blackford, J., & Patel, S. (2019). Endocannabinoid signaling collapse mediates stress-induced amygdalo-cortical strengthening. Neuron, S0896-6273(19)31090–6. https://doi.org/10.1016/j.neuron.2019.12.024

43. Gass, J. T., & Chandler, L. J. (2013). The plasticity of extinction: contribution of the prefrontal cortex in treating addiction through inhibitory learning. Frontiers in Psychiatry, 4(46). https://doi.org/10.3389/fpsyt.2013.00046

44. Jiang, C., Wang, X., Le, Q., Liu, P., Liu, C., Wang, Z., He, G., Zheng, P., Wang, F., & Ma, L. (2019). Morphine coordinates SST and PV interneurons in the prelimbic cortex to disinhibit pyramidal neurons and enhance reward. Molecular Psychiatry. https://doi.org/10.1038/s41380-019-0480-7

45. Pati, D., Pina, M. M., & Kash, T. L. (2019). Ethanol-induced conditioned place preference and aversion differentially alter plasticity in the bed nucleus of stria terminalis. Neuropsychopharmacology, 44(11), 1843–1854. https://doi.org/10.1038/s41386-019-0349-0

46. Varodayan, F. P., De Guglielmo, G., Logrip, M. L., George, O., & Roberto, M. (2017). Alcohol dependence disrupts amygdalar L-type voltage-gated calcium channel mechanisms. Journal of Neuroscience, 37(17), 4593–4603. https://doi.org/10.1523/JNEUROSCI.3721-16.2017

47. Salling, M. C., Skelly, M. J., Avegno, E., Regan, S., Zeric, T., Nichols, E., & Harrison, N. L. (2018). Alcohol consumption during adolescence in a mouse model of binge drinking alters the intrinsic excitability and function of the prefrontal cortex through a reduction in the hyperpolarization-activated cation current. Journal of Neuroscience, 38(27), 6207–6222. https://doi.org/10.1523/JNEUROSCI.0550-18.2018

48. Gu, N., Vervaeke, K., Hu, H., & Storm, J. F. (2005). Kv7/KCNQ/M and HCN/h, but not KCa2/SK channels, contribute to the somatic medium after-hyperpolarization and excitability control in CA1 hippocampal pyramidal cells. Journal of Physiology, 566(3), 689–715. https://doi.org/10.1113/jphysiol.2005.086835

49. Xu, Han, Jeong, H. Y., Tremblay, R., & Rudy, B. (2013). Neocortical Somatostatin-Expressing GABAergic Interneurons Disinhibit the Thalamorecipient Layer 4. Neuron, 77(1), 155–167. https://doi.org/10.1016/j.neuron.2012.11.004

50. Hughes, B. A., Bohnsack, J. P., O’Buckley, T. K., Herman, M. A., & Morrow, A. L. (2019). Chronic ethanol exposure and withdrawal impair synaptic GABAA receptor-mediated neurotransmission in deep-layer prefrontal cortex. Alcoholism: Clinical and Experimental Research, 43(5), 822–832. https://doi.org/10.1111/acer.14015

51. Gao, M., Liu, C. L., Yang, S., Jin, G. Z., Bunney, B. S., & Shi, W. X. (2007). Functional coupling between the prefrontal cortex and dopamine neurons in the ventral tegmental area. Journal of Neuroscience, 27(20), 5414–5421. https://doi.org/10.1523/JNEUROSCI.5347-06.2007

52. McGarry, L. M., & Carter, A. G. (2017). Prefrontal cortex drives distinct projection neurons in the basolateral amygdala. Cell Reports, 21(6), 1426–1433. https://doi.org/10.1016/j.celrep.2017.10.046

53. Siciliano, C. A., Noamany, H., Chang, C. J., Brown, A. R., Chen, X., Leible, D., Lee, J. J., Wang, J., Vernon, A. N., Vander Weele, C. M., Kimchi, E. Y., Heiman, M., & Tye, K. M. (2019). A cortical-brainstem circuit predicts and governs compulsive alcohol drinking. Science, 366(6468), 1008–1012. https://doi.org/10.1126/science.aay1186

